# Detailed phenotyping of *Tbr1-2A-CreER* knock-in mice demonstrates significant impacts on TBR1 protein levels and axon development

**DOI:** 10.1101/2024.04.04.588147

**Authors:** Marissa Co, Grace K. O’Brien, Kevin M. Wright, Brian J. O’Roak

## Abstract

Spatiotemporal control of Cre-mediated recombination has been an invaluable tool for understanding key developmental processes. For example, knock-in of *Cre* into cell type marker gene loci drives *Cre* expression under endogenous promoter and enhancer sequences, greatly facilitating the study of diverse neuronal subtypes in the cerebral cortex. However, insertion of exogenous DNA into the genome can have unintended effects on local gene regulation or protein function that must be carefully considered. Here, we analyze a recently generated *Tbr1-2A-CreER* knock-in mouse line, where a *2A-CreER* cassette was inserted in-frame just before the stop codon of the transcription factor gene *Tbr1*. Heterozygous *TBR1* mutations in humans and mice are known to cause autism or autism-like behavioral phenotypes accompanied by structural brain malformations, most frequently a reduction of the anterior commissure. Thus, it is critical for modified versions of *Tbr1* to exhibit true wild-type-like activity. We evaluated the *Tbr1-2A-CreER* allele for its potential impact on *Tbr1* function and complementation to *Tbr1* loss-of-function alleles. In mice with one copy of the *Tbr1-2A-CreER* allele, we identified reduction of TBR1 protein in early postnatal cortex along with thinning of the anterior commissure, suggesting hypersensitivity of this structure to TBR1 dosage. Comparing *Tbr1-2A-CreER* and *Tbr1*-null heterozygous and homozygous mice to *Tbr1*-null complementation crosses showed reductions of TBR1 dosage ranging from 28.4% to 95.9%. Using these combinatorial genotypes, we found that low levels of TBR1 protein (∼16%) are sufficient to establish cortical layer positioning, while greater levels (>50%) are required for normal suppression of layer 5 identity. In total, these results strongly support the conclusion that *Tbr1-2A-CreER* is a hypomorphic allele. We advise caution when interpreting experiments using this allele, such as transcriptomic studies, considering the sensitivity of various corticogenic processes to TBR1 dosage and the association of heterozygous *TBR1* mutations with complex neurodevelopmental disorders.

## INTRODUCTION

The engineering of transgenic mice expressing site-specific recombinases under specific gene regulatory elements has permitted experimental access to distinct cellular subpopulations across tissues and developmental stages. In the widely used Cre-lox system, Cre catalyzes the recombination of DNA between two loxP recognition sequences. Thus, breeding *Cre* driver mouse lines with loxP-carrying lines can allow for cell type-specific gene knockouts or transgene activation/deactivation (Lakso et al., 1992; Orban et al., 1992; Gu et al., 1994). Furthermore, Cre activity in mice can be temporally controlled in a dose-dependent manner through fusion of Cre to a mutated domain of the estrogen receptor (CreER), which becomes active with the administration of tamoxifen (Metzger and Chambon, 2001). To achieve cell type-specificity of *Cre* expression in mice, one widely used approach is targeted knock-in of *Cre* into an endogenous gene locus using homologous recombination (Jin et al., 2000; Lomelí et al., 2000). Building upon this strategy, inclusion of additional elements can allow for bicistronic expression of both the targeted gene and *Cre*, which theoretically leaves expression of the targeted gene intact. One such element is an internal ribosome entry site (IRES), which allows ribosomes to independently initiate translation of two distinct proteins from a single *‘gene-IRES-Cre’* transcript (Stanley et al., 2002). Alternatively, 2A sequences permit translation of two proteins from a single ‘*gene-2A-Cre’* transcript through ribosomal “skipping” of glycyl-prolyl peptide bond formation during 2A translation (Donnelly et al., 2001; Engert et al., 2009). This mechanism adds extra amino acids to the C-terminus of the polypeptide upstream of 2A, while an extra proline is added to the N-terminus of the downstream polypeptide.

In the field of neuroscience, numerous Cre driver mouse lines have been generated to study neuronal diversity in the cerebral cortex, which is organized into functional areas and cytoarchitectural layers (Gerfen et al., 2013; Harris et al., 2014; Daigle et al., 2018; Cadwell et al., 2019; Matho et al., 2021). The majority of cortical neurons (∼80%) are glutamatergic projection neurons (PNs) that exhibit substantial diversity in their transcriptomic profiles, connectivity, and electrophysiological properties across areas and layers (Saunders et al., 2018; Tasic et al., 2018; Yao et al., 2021). Several Cre lines used for studying cortical PNs express Cre under the promoter of a transcription factor (TF) gene, taking advantage of subtype-specific expression patterns of TFs that typically align with their biological roles in driving subtype identity. Examples of Cre lines harnessing subtype-specific TF expression patterns include *Cux2-Cre* (L2-4 intratelencephalic PNs), *Tlx3-Cre* (L5 intratelencephalic PNs), *Fezf2-2A-CreER* (L5 pyramidal tract PNs), and *Foxp2-IRES-Cre* (L6 corticothalamic PNs) (Franco et al., 2011; Gerfen et al., 2013; Rousso et al., 2016; Matho et al., 2021). Importantly, over the past decade, heterozygous mutations in many of these same TF genes have been identified as highly penetrant risk factors for complex neurodevelopmental disorders, such as autism, intellectual disability, epilepsy, and language disorders (Chatron et al., 2018; Feliciano et al., 2019; Satterstrom et al., 2020).

While bicistronic Cre driver knock-in lines typically preserve expression of the targeted gene, unintended effects can occur. Insertion of exogenous DNA sequences into the genome can produce off-target effects on proximal or even distal gene expression (Olson et al., 1996; Pham et al., 1996; Meier et al., 2010). Additionally, the mechanism of 2A ribosomal skipping appends extra amino acids to the termini of each 2A-linked protein, which in rare cases can interfere with protein stability or function (Lengler et al., 2005; Hasegawa et al., 2007; Reinhardt et al., 2020). Moreover, ribosome read-through at the 2A sequence can produce fusion proteins of unpredictable function (Liu et al., 2017; Velychko et al., 2019). Thus, it is imperative to validate proper expression and function of targeted genes in Cre driver knock-in lines and to document any unexpected effects.

Here, we analyzed a recently generated *Tbr1-2A-CreER* line to validate *Tbr1* expression during early postnatal cortical development (Matho et al., 2021). *Tbr1* encodes a transcription factor restricted to specific neuronal subtypes during cortical development, including early-born deep-layer glutamatergic PNs and Cajal-Retzius cells (Hevner et al., 2001). In humans, heterozygous *de novo* mutations in *TBR1* lead to a complex neurodevelopmental disorder that commonly includes autism, intellectual disability, behavioral disturbances, and speech and motor delays (O’Roak et al., 2012b; O’Roak et al., 2012a; O’Roak et al., 2014; Nambot et al., 2020). Moreover, *TBR1* mutations lead to altered cortical development in both humans and mice (Hevner et al., 2001; Huang et al., 2014; Vegas et al., 2018; Nambot et al., 2020; Co et al., 2022). We found that the *Tbr1-2A-CreER* allele causes decreased TBR1 protein levels in early postnatal cortex. Furthermore, through classical genetic complementation experiments, we leveraged a series of *Tbr1* allele combinations to assess the sensitivity of different corticogenic processes to TBR1 dosage.

## MATERIALS AND METHODS

### Animals

All animal procedures were approved by the Oregon Health & Science University Institutional Animal Care and Use Committee. All mice were maintained on a C57BL/6NJ background (The Jackson Laboratory, strain #005304). *Tbr1-*null mice were generated by germline deletion using *Sox2-Cre* of the *Tbr1* conditional allele with floxed exons 2-3 (provided by J. Rubenstein) (Fazel Darbandi et al., 2018). *Tbr1-2A-CreER* mice were obtained from The Jackson Laboratory (strain #036299) (Matho et al., 2021). *Tbr1*-null and *Tbr1-2A-CreER* mice were PCR genotyped using primers in **Supplementary Table 1**. In-frame insertion of the *2A-CreER* cassette was verified by Sanger sequencing using the primer in **Supplementary Table 1**. The following crosses were used to generate experimental animals: (1) *Tbr1*^*+/creER*^ × *Tbr1*^*+/creER*^, (2) *Tbr1*^*+/creER*^ × *Tbr1*^*+/–*^, (3) *Tbr1*^*+/–*^ × *Tbr1*^*+/–*^. Mice were group housed under a 12 h light/dark cycle and given *ad libitum* access to food and water.

### Experimental Design and Statistical Analysis

Postnatal day 0 (P0) mice of each sex were used for all experiments. Cohorts for each experiment were composed of littermate wild-type (WT) and *Tbr1* variant mice from at least two independent litters. Data plotting and statistical tests were performed using Prism 10 (GraphPad Software). Data are represented as the mean ± SEM. Each dot represents one animal. Analyses across groups were performed using one-way ANOVA with Tukey’s multiple-comparisons test. Significance was defined as *p* < 0.05.

### Western Blot

Whole cortex was dissected at P0, flash frozen in liquid nitrogen, and stored at –80°C until all samples were collected. Frozen tissue from one cortical hemisphere per mouse (∼30 mg at P0) was dounce homogenized in ice-cold RIPA buffer (150 mM NaCl, 1% IGEPAL CA-630, 0.5% sodium deoxycholate, 0.1% SDS, 50 mM Tris-HCl pH 8.0, 2mM EDTA pH 8.0) containing cOmplete™ Protease Inhibitor Cocktail (Roche). Nuclei were lysed using a probe sonicator (Misonix, model XL-2000) at setting 6 with 10 s ON / 20 s OFF intervals for two rounds. Lysates were incubated on ice for 30 min, then centrifuged at 10,000 *rcf* for 10 min at 4°C to remove debris. Protein concentrations were determined using the Pierce BCA Protein Assay Kit (Thermo Fisher Scientific). Lysates were boiled in 1× Laemmli SDS sample buffer (Thermo Fisher Scientific) at 95°C for 5 min and stored at –20°C until SDS-PAGE. Total protein (30 μg/sample) was resolved on 4–15% polyacrylamide gels (Bio-Rad) and transferred to PVDF membranes. Membranes were incubated in block solution (5% milk in TBS with 0.1% Tween-20 [TBST]) for 1 h at room temperature (RT), incubated in primary antibodies in block solution for 24–72 h at 4°C, washed in TBST 4 times for 5 min each, incubated in secondary antibodies in block solution for 1 h RT, washed in TBST 4 times for 5 min each, and imaged using an Odyssey CLx with Image Studio software (LI-COR). The following primary antibodies and dilutions were used: rabbit anti-TBR1 (1:1000, Abcam, catalog #ab31940); rabbit anti-Cre Recombinase (1:1000, BioLegend, catalog #PRB-106P); mouse anti-GAPDH (1:5000, Proteintech, catalog #60004-1-Ig). The following secondary antibodies and dilutions were used: IRDye® 800CW Donkey anti-Rabbit IgG (1:15,000, LI-COR Biosciences, catalog #926-32213); IRDye® 680RD Donkey anti-Mouse IgG (1:15,000, LI-COR Biosciences, catalog #926-68072). Western blot band signals were measured using Image Studio software. Within each blot, each TBR1 or Cre band signal was normalized to its corresponding GAPDH loading control signal, and then each normalized signal was adjusted to the average normalized signal across control replicates.

### Reverse Transcription Quantitative PCR (RT-qPCR)

Cortex was dissected at P0, flash frozen in liquid nitrogen, and stored at –80°C until all samples were collected. Total RNA was extracted using the RNeasy Mini Kit (QIAGEN, catalog #74104) according to the manufacturer protocol. Tissue was lysed by trituration in Buffer RLW followed by centrifugation through QIAshredder columns (QIAGEN, catalog #79654). On-column DNase digestion was performed using RNase-Free DNase Set (QIAGEN, catalog #79254) according to the manufacturer protocol. cDNA was synthesized from 1 μg total RNA using the ProtoScript II First Strand cDNA Synthesis Kit (New England Biolabs, catalog #E6560S) with oligo-dT priming according to the manufacturer protocol. cDNA templates and no RT controls were diluted 1:20 for multiplexed PrimeTime qPCR Assays (Integrated DNA Technologies). Primer and probe sequences are provided in **Supplementary Table 1**. qPCR assays were run in triplicate. Probe fluorescence was measured with a QuantStudio 5 Real-Time PCR System (Thermo Fisher Scientific) running the following cycling program: 95°C for 3 min, 40 cycles of 95°C for 15 s and 60°C for 1 min, and 4°C hold. qPCR primer efficiencies were measured using WT cortex cDNA (*Tbr1, Actb*) or *Tbr1*^*+/creER*^ cortex cDNA (*Cre*) for the standard curve and fell within 90–110%. After averaging across technical replicates for each sample, *Tbr1* or *Cre* fold gene expression was calculated using the 2^−Δ ΔCt^ method with *Actb* as the reference gene. Each sample was adjusted to the control genotype average within its respective experimental batch.

### Immunohistochemistry (IHC)

P0 whole brains were drop fixed in 4% EM-grade paraformaldehyde (PFA) overnight at 4°C, then rinsed in PBS. Brains were embedded in 3% low-melting point (LMP) agarose and sectioned coronally at 100 μm using a vibratome (Leica, model VT1200). Free-floating sections were incubated in block solution (2% donkey serum, 0.2% Triton X-100 in PBS) for 30 min RT, then incubated in primary antibodies in block solution for 48–72 h at 4°C. After washes in PBS for >5 h RT, sections were incubated in secondary antibodies and Hoechst 33342 (1:5000, Thermo Fisher Scientific, catalog #H3570) in block solution overnight RT. After washes in PBS for >5 h RT, sections were mounted onto glass slides and coverslipped with Fluoromount-G (Southern Biotech). The following primary antibodies and dilutions were used: rabbit anti-CDP/CUX1 (1:500, Santa Cruz Biotechnology, catalog #sc-13024); rat anti-CTIP2 (1:500, Abcam, catalog #ab18465); rat anti-L1 (1:500, Millipore, catalog #MAB5272); rabbit anti-TBR1 (1:500, Millipore, catalog #AB10554). The following secondary antibodies and dilutions were used: Donkey anti-Rabbit IgG (H+L) Alexa Fluor 488 (1:500, Thermo Fisher Scientific, catalog #A-21206), Donkey anti-Rat IgG (H+L) Alexa Fluor 546 (1:500, Immunological Sciences, catalog #IS20320).

## RESULTS

### Mice with *Tbr1-2A-CreER* allele show reduced TBR1 protein in cortex

As previously described, Matho and colleagues generated the *Tbr1-2A-CreER* allele by inserting a *2A-CreER* cassette in-frame before the stop codon of *Tbr1* (**Fig. 1A**) (Matho et al., 2021). We verified in-frame insertion of the cassette by Sanger sequencing of genomic DNA from homozygous *Tbr1-2A-CreER* mice (data not shown). To assess the impacts of the *2A-CreER* cassette on *Tbr1* function, we analyzed heterozygous and homozygous *Tbr1-2A-CreER* mice (i.e., *Tbr1*^*+/creER*^, *Tbr1*^*creER/creER*^) as well as offspring from *Tbr1*^*+/creER*^ × *Tbr1*^*+/–*^ complementation crosses (i.e., *Tbr1*^*+/creER*^, *Tbr1*^*+/–*^, *Tbr1*^*creER/–*^) and *Tbr1*^*+/–*^ × *Tbr1*^*+/–*^ crosses. All genotypes were directly compared to their *Tbr1*^*+/+*^ littermate controls. If the *2A-CreER* cassette had no impact on *Tbr1* function, we would expect *Tbr1*^*+/creER*^ and *Tbr1*^*creER/creER*^ mice to be phenotypically indistinguishable from *Tbr1*^*+/+*^ mice. Additionally, *Tbr1*^*creER/–*^ mice would phenocopy *Tbr1*^*+/–*^ mice, which have known reductions in TBR1 protein in cortex and thinning of the posterior limb of the anterior commissure (AC) (Bulfone et al., 1998; Huang et al., 2014; Co et al., 2022).

**Figure 1.**
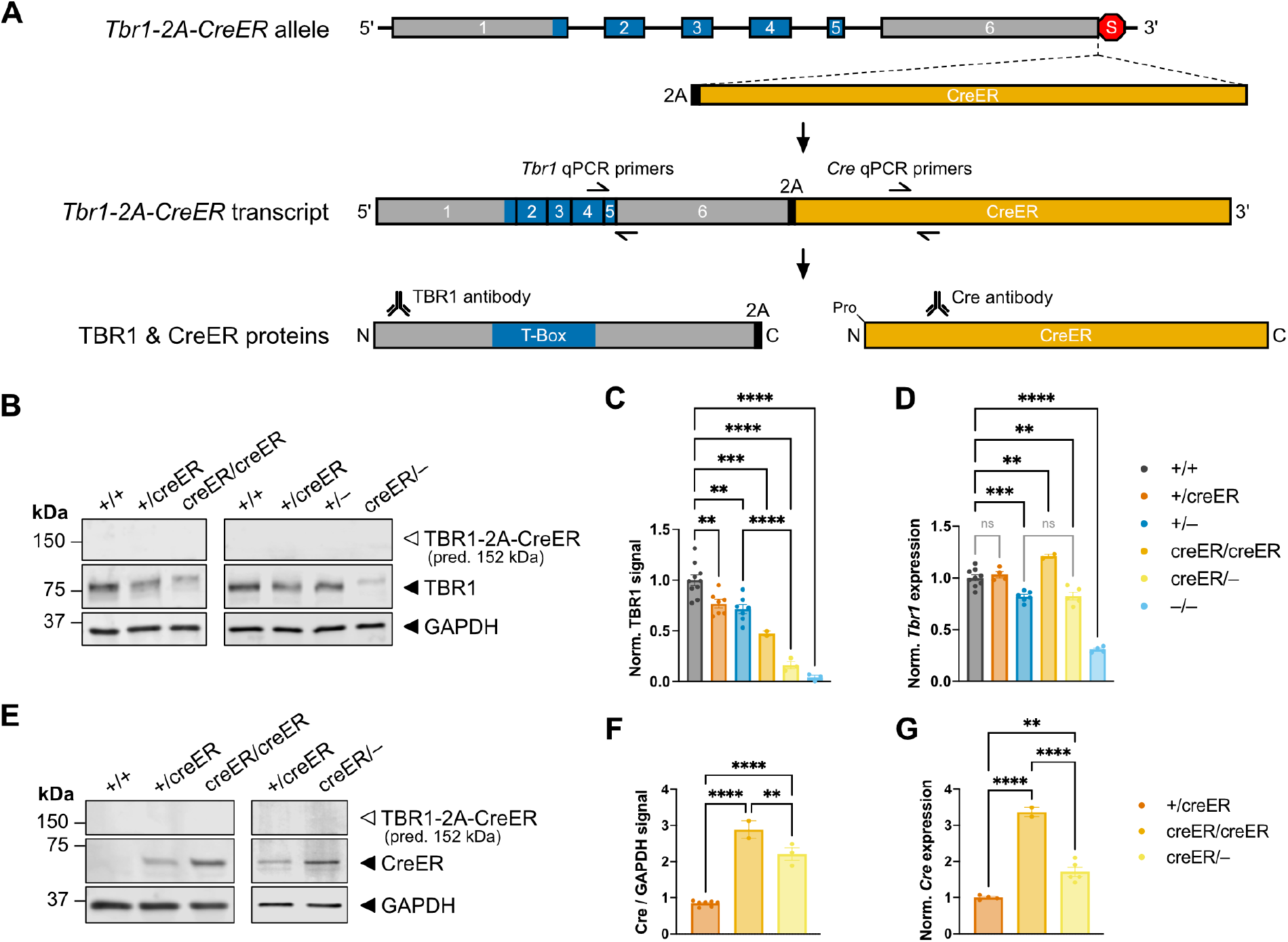
Knock-in of *2A-CreER* at the 3’ end of *Tbr1* reduces TBR1 protein levels in postnatal cortex. (***A***) *Tbr1-2A-CreER* mechanism for driving CreER in *Tbr1*+ cells. A *2A-CreER* cassette was inserted in-frame before the stop codon of mouse *Tbr1* (Matho et al., 2021). Regions in blue encode the T-Box DNA-binding domain. Short linkers (9-12 bp) are present between *Tbr1, 2A*, and *CreER* sequences (not shown). A single transcript generated from the allele produces independent TBR1 and CreER proteins through ribosome skipping at 2A. Twenty amino acids from the linker and 2A are added to the C-terminus of TBR1, while an extra proline (Pro) is added to the N-terminus of CreER. Approximate primer and antibody locations for qPCR and Western blot, respectively, are indicated. (***B-C***) Western blots for TBR1 from cortical lysates of P0 *Tbr1* mutant mice and WT littermates (n = 2-10 mice/genotype). GAPDH was used as loading control. 2A-tagged TBR1 migrates at higher molecular weight in *Tbr1*^*creER/creER*^ and *Tbr1*^*creER/–*^. Predicted molecular weight of TBR1-2A-CreER fusion protein is indicated (open arrowhead), but this protein is not detected. (***D***) RT-qPCR for *Tbr1* from cortical cDNA of P0 *Tbr1* mutant mice and WT littermates (n = 2-10 mice/genotype). (***E-F***) Western blots for CreER from cortical lysates of P0 *Tbr1* mutant mice and WT littermates (n = 2-7 mice/genotype). GAPDH was used as loading control. (***G***) RT-qPCR for *Cre* from cortical cDNA of P0 *Tbr1* mutant mice and WT littermates (n = 2-7 mice/genotype). Data are plotted as mean ± SEM. Each dot represents one animal. One-way ANOVA with Tukey’s multiple comparisons test was used in ***C, D, F, G***. ^**^p < 0.01; ^***^p < 0.001; ^****^p < 0.0001; ns: not significant.

We first examined TBR1 protein levels by Western blot of whole cortical lysates from P0 mice. We observed significantly reduced TBR1 signal in each experimental genotype compared to *Tbr1*^*+/+*^, with reductions of 23.3% (*Tbr1*^*+/creER*^), 28.4% (*Tbr1*^*+/–*^), 52.5% (*Tbr1*^*creER/creER*^), 83.8% (*Tbr1*^*creER/–*^), and 95.9% (*Tbr1*^*–/–*^) (**Fig. 1B-C**). In *Tbr1*^*creER/creER*^ and *Tbr1*^*creER/–*^, we observed a higher molecular weight band corresponding to 2A-tagged TBR1 (**Fig. 1B**), which is predicted to have 20 additional amino acids at its C-terminus based on our Sanger sequencing data. In all *CreER*+ genotypes, we did not observe bands corresponding to a putative TBR1-2A-CreER fusion protein (predicted size 152 kDa) using either TBR1 or Cre antibodies (**Fig. 1B, 1E**), indicating that ribosome read-through does not account for reduced TBR1. When blotting with α-Cre across genotypes, we noted that CreER levels did not follow the near-linear decrease observed for TBR1 (**Fig. 1C, 1F**). To distinguish whether TBR1 reductions were occurring at the transcript or protein level, we performed RT-qPCR in P0 cortex for *Tbr1* and *CreER*. Only the genotypes with a *Tbr1*-null allele (i.e., *Tbr1*^*+/–*^, *Tbr1*^*creER/–*^, *Tbr1*^*–/–*^) showed significant reductions in *Tbr1* transcript (**Fig. 1D**). Moreover, *CreER* transcript levels were consistent with CreER protein levels, while *Tbr1* transcript levels were not consistent with TBR1 protein levels (**Fig. 1C-D, 1F-G**). These findings suggest reduction of TBR1 at the protein level in *Tbr1-2A-CreER* mice. Of note, interpretation of TBR1 and CreER protein versus transcript levels in *Tbr1* mutant mice is complicated by TBR1’s potential negative autoregulatory functions (Co et al., 2022). Nonetheless, these results demonstrate that the *Tbr1-2A-CreER* allele leads to decreased TBR1 protein in postnatal cortex.

### Altered olfactory bulb and cortical development in mice with low TBR1 dosage

*Tbr1*^*–/–*^ mice show severe defects in postnatal survival (death ∼P0-P3), olfactory bulb size, neocortical lamination, neuronal subtype specification, and axon tract formation (Bulfone et al., 1998; Hevner et al., 2001; Han et al., 2011; McKenna et al., 2011). In contrast, *Tbr1*^*+/–*^ mice survive to adulthood with slightly reduced olfactory bulb size, grossly normal neocortical formation, and selective reduction of AC posterior limb axons (Huang et al., 2014; Huang et al., 2019; Co et al., 2022). We utilized our genotypic series of *Tbr1-2A-CreER* mutants, ranging from ∼23% to ∼96% TBR1 reduction, to assess TBR1 dosage thresholds for these developmental phenotypes. *Tbr1*^*+/creER*^ and *Tbr1*^*creER/creER*^ mice are healthy and viable, although *Tbr1*^*creER/creER*^ breeders are reported by The Jackson Laboratory to show high rates of nonproductivity. *Tbr1*^*creER/–*^ mice survive longer than *Tbr1*^*–/–*^ but show growth delays by P7, leading to humane euthanasia by ∼P18 due to low body weight (data not shown). Examination of *Tbr1*^*creER/–*^ brains at P0 revealed overtly normal brain morphology but severely reduced olfactory bulb size, similar to *Tbr1*^*–/–*^ (**Fig. 2A**) (Hevner et al., 2001).

**Figure 2.**
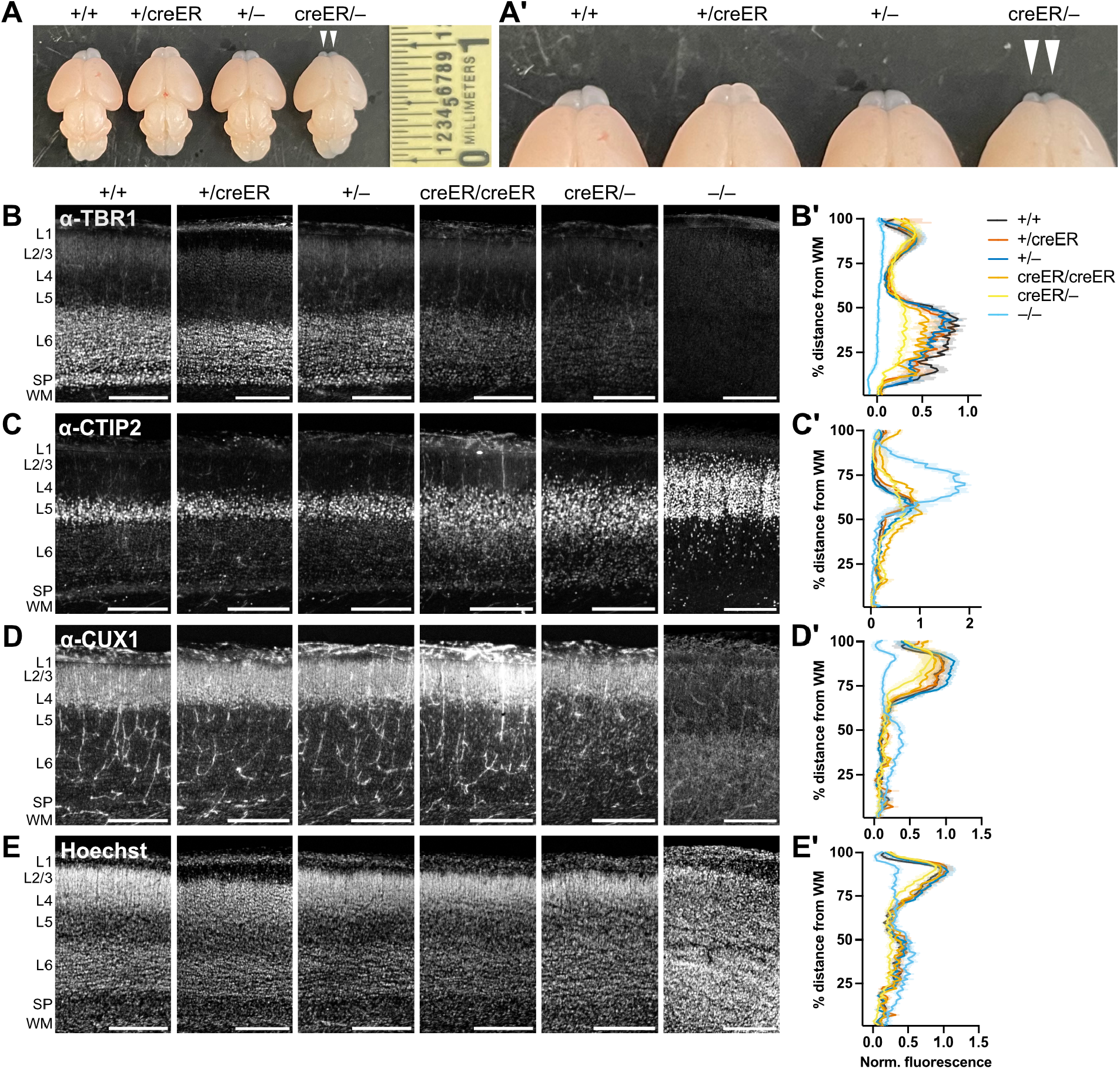
TBR1 dosage effects on olfactory bulb development and neocortical layering. (***A, A’***) P0 brains from *Tbr1* mutant mice and WT littermate. Arrowheads show reduced olfactory bulb size in *Tbr1*^*creER/–*^ mouse. (***B-E***) IHC and fluorescence intensity for TBR1 (***B, B’***), L5 marker CTIP2 (***C, C’***), L2-4 marker CUX1 (***D, D’***), and nuclear marker Hoechst (***E, E’***) in P0 cortex (n = 2-6 mice/genotype). L: layer; SP: subplate; WM: white matter. Scale bars: ***B, C, D, E***, 200 μm. Data are plotted as mean ± SEM.

We performed immunohistochemistry (IHC) to examine neocortical lamination and neuron subtype specification in *Tbr1-2A-CreER* mice at P0, comparing them to *Tbr1*^*+/–*^ and *Tbr1*^*–/–*^ mutants. IHC for TBR1 showed a graded reduction of TBR1 signal in layer (L) 6 of primary somatosensory cortex across mutant genotypes (**Fig. 2B**), reflecting the TBR1 reductions observed by Western blot analysis (**Fig. 1B-C**). IHC for TBR1, CTIP2 (L5 marker), and CUX1 (L2-4 marker) indicated normal deep-layer vs. upper-layer positioning in all genotypic combinations containing the *Tbr1-2A-CreER* allele (**Fig. 2C-D**), suggesting that the ∼16% of normal TBR1 levels in *Tbr1*^*creER/–*^ cortex is sufficient for normal positioning of layers (**Fig. 1C**). This result contrasted with the inverted CTIP2+ and CUX1+ layer positions in *Tbr1*^*–/–*^ cortex, in which we observed only ∼4% of WT TBR1 levels (**Fig. 1C, Fig. 2C-D**). In *Tbr1*^*creER/creER*^ and *Tbr1*^*creER/–*^ cortex, which express <50% of WT TBR1 levels (**Fig. 1C**), we observed increased CTIP2+ neurons and expansion of L5 width, both above and below the normal relative bounds of L5 (**Fig. 2C**, orange and yellow traces). L6 and L5 were also less distinguishable by Hoechst nuclear staining in *Tbr1*^*creER/–*^ neocortex (**Fig. 2E**). These findings suggest impaired L6 neuronal specification and migration in *Tbr1*^*creER/creER*^ and *Tbr1*^*creER/–*^ neocortex, as L6 neurons acquire L5-like properties in the absence of *Tbr1* (Han et al., 2011; McKenna et al., 2011; Fazel Darbandi et al., 2018). We also observed decreased relative width of the upper layers in *Tbr1*^*creER/–*^ neocortex, based on CUX1 IHC and Hoechst staining (**Fig. 2D-E**). Together, these results further demonstrate that the *Tbr1-2A-CreER* allele is not functionally equivalent to WT. They also indicate that ∼16% of normal TBR1 level in developing neocortex is sufficient to establish overall layer positioning, but <50% is insufficient to fully suppress L5 marker expression in deep-layer neurons.

### Development of the anterior commissure is highly sensitive to TBR1 dosage

Formation of the AC is known to be impacted by disruption of one copy of *TBR1* in both humans and mice, across independent mutant alleles (Huang et al., 2014; Nambot et al., 2020; Co et al., 2022). We performed IHC for the axon marker L1 in P0 *Tbr1-2A-CreER* mutants to examine how TBR1 dosage impacts the development of the AC, as well as the corpus callosum (**Fig. 3A**). We observed thinning of the AC posterior limb in *Tbr1*^*+/creER*^ mice (N = 7 of 7 showing thinned posterior limb), indicating that one copy of the *Tbr1-2A-CreER* allele is sufficient to impact this axon tract (**Fig. 3B**, arrowheads). Similar to *Tbr1*^*+/–*^, the AC posterior limb was severely reduced or absent in *Tbr1*^*creER/creER*^ mice (N = 3) and *Tbr1*^*creER/–*^ mice (N = 5) (**Fig. 3C-E**, arrowheads). The AC anterior limb was also reduced in *Tbr1*^*creER/creER*^ and *Tbr1*^*creER/–*^ mice (**Fig. 3C-E**, arrowheads). We observed increased frequency of axon pathfinding errors in *Tbr1*^*creER/creER*^ and *Tbr1*^*creER/–*^ mice (**Fig. 3D**, white arrow, for example). Furthermore, we identified callosal Probst bundles in *Tbr1*^*creER/–*^ mice, similar to *Tbr1*^*–/–*^ (**Fig. 3E-F**, yellow arrows). Other severe axon phenotypes were unique to *Tbr1*^*–/–*^ mice, including absence of both anterior and posterior AC limbs (**Fig. 3F**, arrowheads) and presence of ectopic cortical axons (**Fig. 3F**, asterisks). Overall, these results indicate that axon developmental processes throughout the brain have differing sensitivities to TBR1 dosage, with the AC posterior limb being highly sensitive to even low-level changes in TBR1.

**Figure 3.**
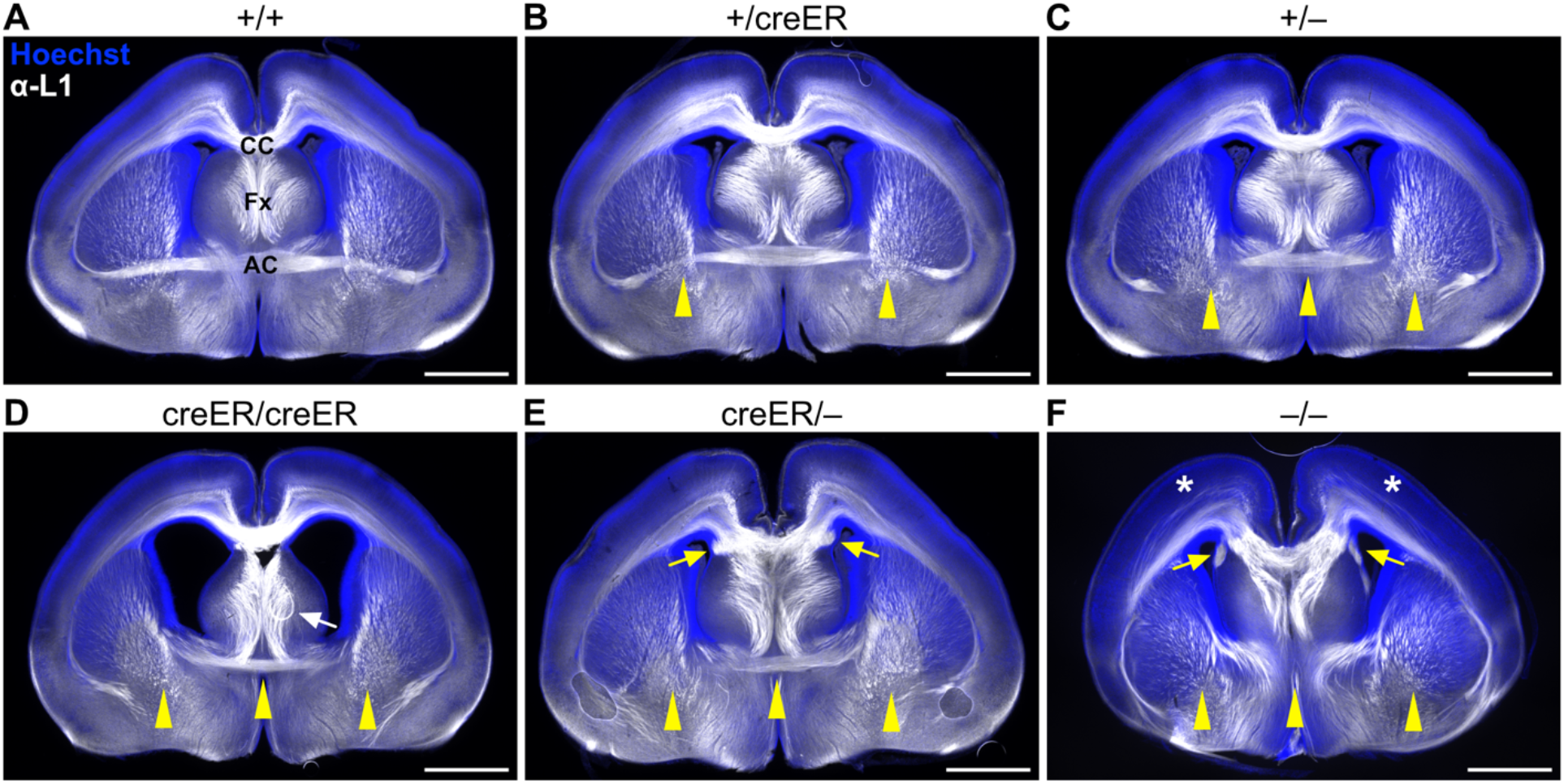
TBR1 dosage effects on callosal and anterior commissure axon development. (***A-F***) IHC for axon marker L1 (white) and Hoechst nuclear stain (blue) in P0 coronal brain sections. Yellow arrowheads indicate thinning or absence of AC in ***B, C, D, E***. White arrow indicates axon pathfinding error in fornix in ***D***. Yellow arrows indicate Probst bundles in corpus callosum of ***E, F***. Asterisks indicate ectopic cortical axons in ***F***. AC: anterior commissure; CC: corpus callosum; Fx: fornix. Scale bars: ***A-F***, 1 mm.

## DISCUSSION

In this study, we identified reduced TBR1 protein and thinning of the AC posterior limb in mice with one copy of the *Tbr1-2A-CreER* allele (Matho et al., 2021). These findings, in comparison with known null alleles, strongly suggest that the integration of the *2A-CreER* cassette into the *Tbr1* locus generated a hypomorphic allele. The AC posterior limb contains axons originating from piriform and entorhinal cortices as well as lateral/temporal portions of neocortex (Fenlon et al., 2021), suggesting PNs in these regions likely exhibit molecular, morphological, and possibly functional abnormalities in *Tbr1*^*+/creER*^ mice. Hence, we advise caution when using this mouse line for analysis of cortical PNs and other cell types where *Tbr1* is expressed. While we did not observe gross neocortical layering defects in *Tbr1*^*+/creER*^ mice, we also did not rule out abnormalities in dendrite development, synapse formation, or synaptic function of deep-layer PNs, which are abnormal in other *Tbr1* mutant mouse models (Fazel Darbandi et al., 2018; Yook et al., 2019; Fazel Darbandi et al., 2020). Human genetic data and animal models indicate that haploinsufficiency mechanisms contribute to *TBR1*-related disorders (OMIM: 60653), pointing to the importance of *TBR1* dosage for normal development (Huang et al., 2014; Landrum et al., 2018; Co et al., 2022). In general, TFs are often dosage-sensitive (Seidman and Seidman, 2002; van der Lee et al., 2020), and thus careful validations must be conducted for Cre driver lines with altered TF loci.

The *Tbr1-2A-CreER* allele does offer advantages for studying the effects of TBR1 dosage on neurodevelopmental processes involving diverse cellular subtypes. For example, we found that TBR1 reductions of up to ∼84% were associated with relatively normal positioning of neocortical layers. Proper layer formation requires Reelin secretion by Cajal-Retzius cells, which express high levels of *Tbr1* and are absent in *Tbr1*^*–/–*^ mice (Hevner et al., 2001; Vilchez-Acosta et al., 2022). In contrast, TBR1 reductions of ∼53% and greater were associated with expansion of CTIP2+ L5 thickness, while TBR1 reduction of ∼23% was associated with thinning of the AC posterior limb. These findings hint at different sensitivities of Cajal-Retzius cells, L6 PNs, and AC-forming PNs to TBR1 protein levels, with the caveat that our molecular analyses were conducted in bulk cortical tissue. Future analyses within specific cellular subtypes may further resolve the mechanisms behind these apparent selective vulnerabilities and may provide insights into the mechanisms driving *TBR1*-associated neurodevelopmental disorders.

## ACKNOWLEDGEMENTS

Research reported in this publication was partially supported by the National Institute of Mental Health of the National Institutes of Health under award number R01MH113926 (B.J.O). The content is solely the responsibility of the authors and does not necessarily represent the official views of the National Institutes of Health. This work was also supported by the OHSU Richard & Marilyn Jones Endowed Fund for Advancement in Autism Research, an award from the OHSU Brain Institute Neuroscience Campaign Fund to Support Brain Health Across the Lifespan (B.J.O.), a Medical Research Foundation of Oregon Early Clinical Investigator Award (M.C.), and a Collins Medical Trust Award (M.C.).

## AUTHOR CONTRIBUTIONS

M.C., K.M.W., and B.J.O. designed experiments; M.C. and G.O. performed experiments; M.C. analyzed data; M.C. wrote the manuscript. All authors edited the manuscript.

